# Urbanisation Reshapes Freshwater Microbiomes: A Systematic Review of Ecological Patterns and Functional Shifts

**DOI:** 10.64898/2026.03.31.715732

**Authors:** Khushi Thakur, Raashi Jain, Shreejit Panda, Hiya Chakma, Aditi Sudhir, Ananya Mukherjee

**Author notes:** **Corresponding Author:** Ananya Mukherjee, School of Arts and Sciences, Azim Premji University Bhopal, Madhya Pradesh, India.

## Abstract

Rapid urbanisation has profoundly shaped microbial diversity across different ecosystems. Freshwater microbiomes are particularly affected by urbanisation activities, such as eutrophication, pollution, runoff, and sewage. This is of significant concern as marginalised communities often depend on waterbodies for their livelihood. Freshwater bodies play a crucial role in maintaining both human and ecological health at population level. Currently, we lack a systematic understanding of the global impacts of urbanisation on freshwater microbiomes in relation to human health, ecosystem functioning, and sustainability.

We identified 90 eligible papers from the last 25 years after screening based on the inclusion exclusion criteria. We extracted data that examined changes in the functional traits such as antimicrobial resistance (AMR), nutrient cycling of the microbiome in urban waterbodies and several other factors. Data were extracted by a thematic analysis followed by a narrative synthesis on specific functional traits. This systematic review presents a comprehensive analysis on the changes and challenges brought about by urbanisation on freshwater bodies. Our results indicate that urbanisation leads to reduced bacterial diversity of urban waterbodies, with a striking increase in reporting of Proteobacteria, Cyanobacteria and Coliform bacteria. These insights will help inform public health strategies and sustainable urban planning.

**Graphical abstract:** 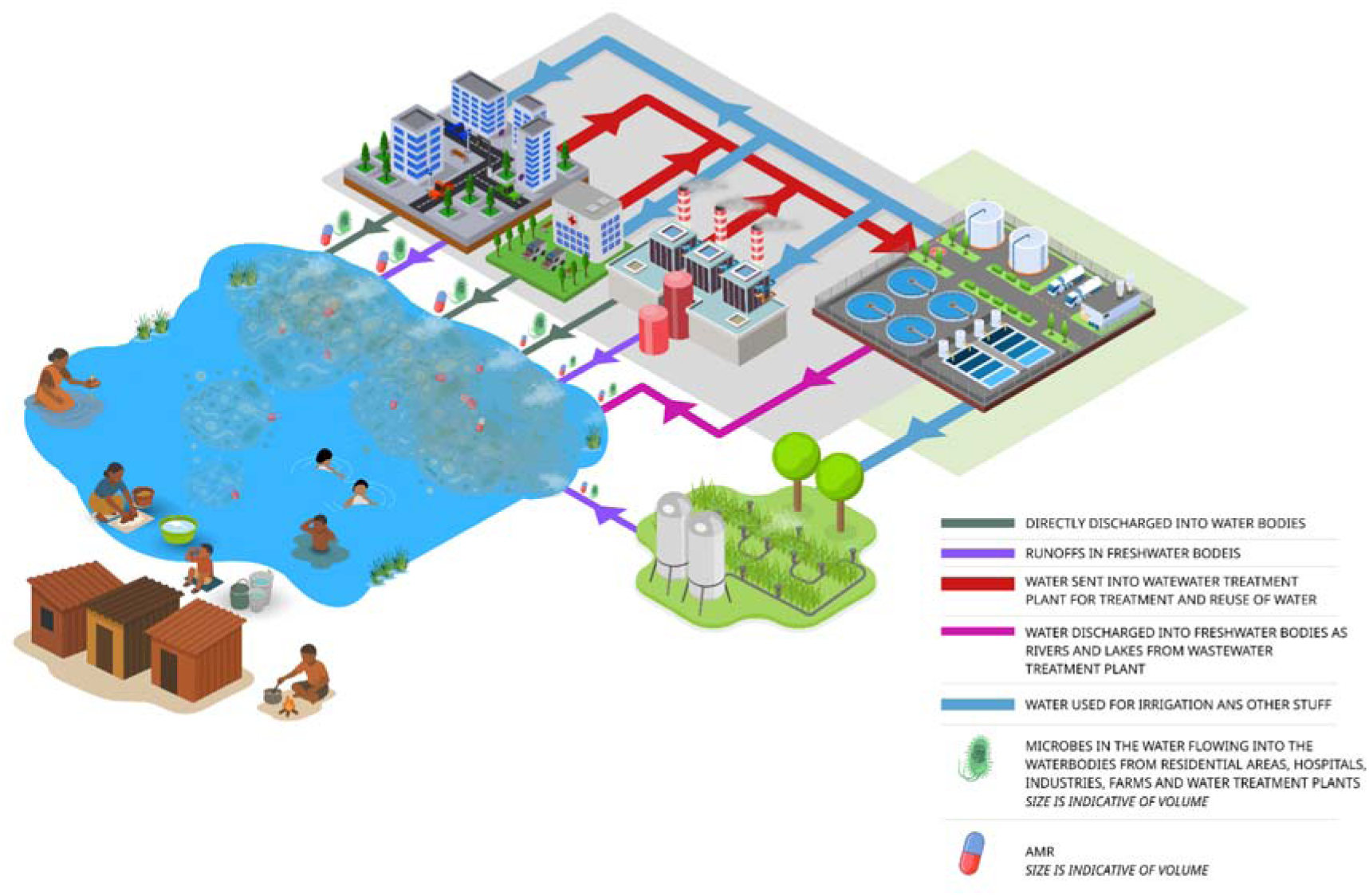

Waterbodies in urban areas function as convergence platforms for anthropogenic and environmental microbiomes. Runoffs, wastewater and effluents contain antimicrobial resistance genes and other pathogens that survive in water due to inadequate treatment. Disposal, use, and overflow of wastewater cause restructuration of microbial communities, proliferation of opportunistic microorganisms, and spread of antimicrobial resistance in aquatic ecosystems.

## Introduction

Cities are the fastest growing ecosystems globally with 81 per cent of the world’s population already residing in urban areas(United Nations, 2025). Urban ecosystems are also highly heterogeneous with a complex mosaic of natural and built environments consisting of buildings, parks, gardens, remnant forests, roads, pavements and waterbodies(Jones et al., 2022). It is no surprise that we live on a planet that is consistently shaped by human activity and cities, the hub of human activity-are rarely designed keeping flora or fauna in mind. Consequently animals often enter cities as their natural habitats are reformed or displaced(Schell et al., 2020). Additionally, microbes, which are fundamental to public health as human commensal microbiota and invisible biodiversity(Matthews et al., 2024), rarely feature in our thinking of urban fauna. This is particularly interesting despite a long history of evidence showing their influence on urban life. John Snow’s seminal work on cholera has shown how the urban dwellers’ interaction with microbes can be very different from those of their rural counterparts(Tulchinsky et al.,2018).

In the last 100 years with the rising use of antibiotics, pollution and the urban rural divide the relationship between humans and microbes has become more complex if not diversified(Kabwe et al., 2020). We are now surrounded by more novel antibiotic resistant genes(ARG) than ever before. Another growing risk is antimicrobial resistance(AMR) as a recent study predicted that in 2050 the global bacterial AMR burden will spell particularly high mortality rates for South Asia, Africa and the Caribbean(Naghavi et al., 2024).

Microbial ecology depicts how ecosystems respond to human-driven climate change. One Health proposes that the well-being of humans is linked to the health(World Health Organization of the ecosystem and the thread that holds it all together are microbial contributions(WHO), 2025). Many studies discuss microbe host associations such as plants, humans and animals, as an eco-holobiont or an extended second genome(Banerjee & Heijden, 2023). Even recently microbial research would heavily focus on pathogens affecting microbial hosts and while understanding zoonotic diseases remain vital, loss of microbial diversity can cause dysbiosis(Berg et al., 2020). Dysbiosis is caused when the composition of microbial diversity drops and leads to the emergence of pathogens which affects the immune system of the host. Thus any dynamic built environment can face the threat of dysbiosis, especially urban landscapes which are constantly shaped by anthropogenic activities. In fact metagenomic maps of urban microbiomes and antimicrobial resistance markers revealed that most cities have a unique microbial fingerprint(Danko et al., 2021).

In this complex mosaic of microbial diversity in urban systems, one crucial and perhaps understudied domain are urban freshwater systems that provide a juxtaposition of ecology and society. Historically habitats and settlements have come up around water bodies whether built or natural such as rivers, lakes, reservoirs, ponds, coastal or inland regions(Cruz-Cano et al., 2025). These continue to underpin urban livelihood, recreational activities, tourism and even play a role in mitigating climate change. They also bear the burden of waste, pollution, anthropogenic activities, toxic waste, fecal contamination and cultural activities. Marginalised communities depend on them for drinking water as well making them directly relevant to public health in the surrounding area(Dickin & Gabrielsson, 2023). This is vital at a time when nearly 4 billion people face water scarcity at least once a year(Mekonnen & Hoekstra, 2016). Thus water management practices become crucial for the health and well being of the population around waterbodies. Urban planning tends to look at waterbodies as passive infrastructure elements instead of thriving ecosystems. Determining the urban waterbody microbiome is not just an exercise in curiosity but data that can assist urban governance and planning, stormwater design, restoration, control pollution and surveil public health.

With the advent of sequencing methods there is no scarcity of microbiome data from specific urban ecosystems, especially waterbodies surrounding a city(González et al., 2025). However, we have a fragmented view of the microbial make-up often in the form of case studies and geographically restricted datasets which prevent us from having a wider context of the ecological patterns around the world. A broader view of the literature can help in looking at both short term fluctuations and long term geographical or climatic trends arising from climate change, pollution or anthropogenic activities. Evidence based urban water management and ecological theory can be strengthened by checking microbial trends beyond mere identification or pathogen dynamics. Fundamental processes such as nutrient cycling and AMR need coordinated comparative approaches.

Keeping this in mind we synthesized insights from 90 studies across three different research search engines to look at the effect urbanisation has on the microbiome of waterbodies in cities across the globe. We have taken a comprehensive approach and shed light on the different human activities observed with respect to the microbial communities observed and nutrient cycling types in global urban freshwater bodies. Our goal was to identify microbes beyond their pathogenic traits and also look at the trend of urbanisation versus the type of microbe identified over time.

## Objectives

The primary questions addressed in this paper are:-

● What evidence exists for the effect of urbanisation on urban freshwater body microbiomes in terms of microbial diversity and functional shifts?
● What are the critical gaps in the available literature and where else can the focus lie?
● How can we look at microbial ecological patterns across geography in urban freshwater bodies?
● Should policy makers focus on microbial make-up of wetlands beyond a few identifying organisms while restoring waterbodies in cities?

This review was conducted to address the objectives. It is following a systematic approach, conforming to the PRISMA reporting standards to ensure comprehensive and unbiased coverage of literature (Page et al., 2020).

## Methods

This study combines systematic review methodology with systematic mapping to identify patterns in microbial taxa reporting, functional traits, and anthropogenic drivers across urban freshwater ecosystems.

We used three different bibliographic databases for this literature review: Web of Science (WOS), Google Scholar (GS) and SCOPUS. Each search string was adapted to suit the database. The search was conducted between January 2025 and March 2025. The number of records retrieved with each string is shown in Table 1. Table 2 contains the rationale for each inclusion and exclusion criteria. We only considered research articles and not reviews.

**Table 1:**
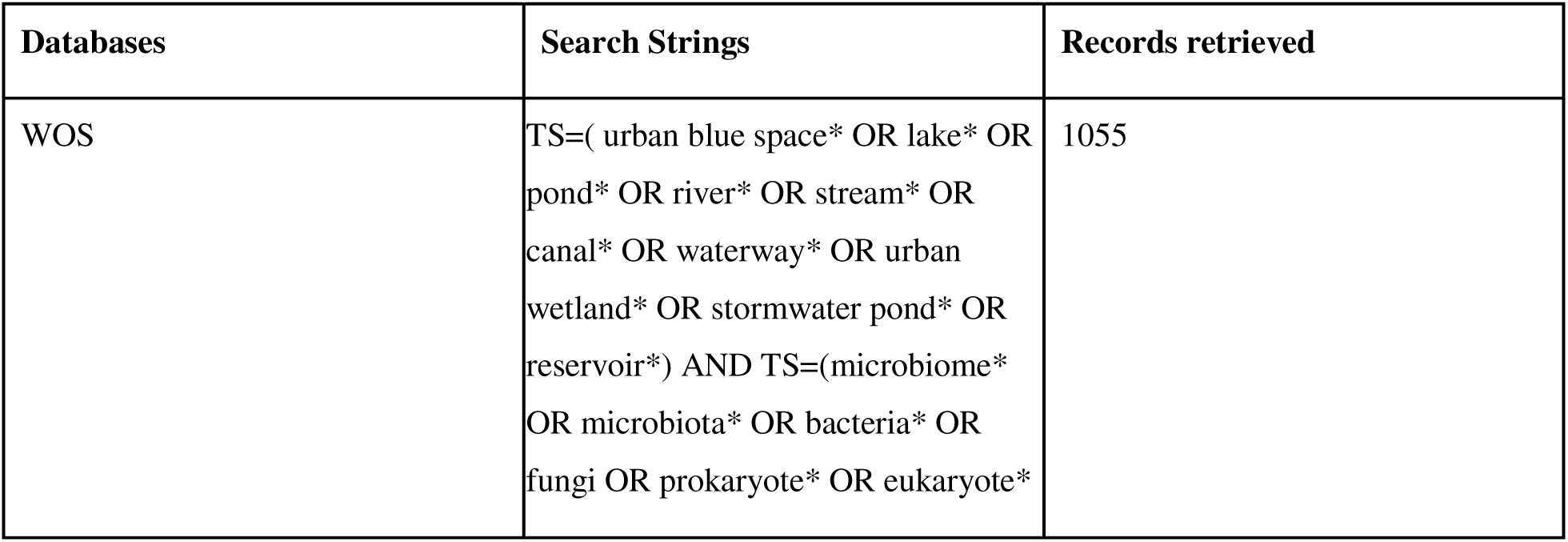
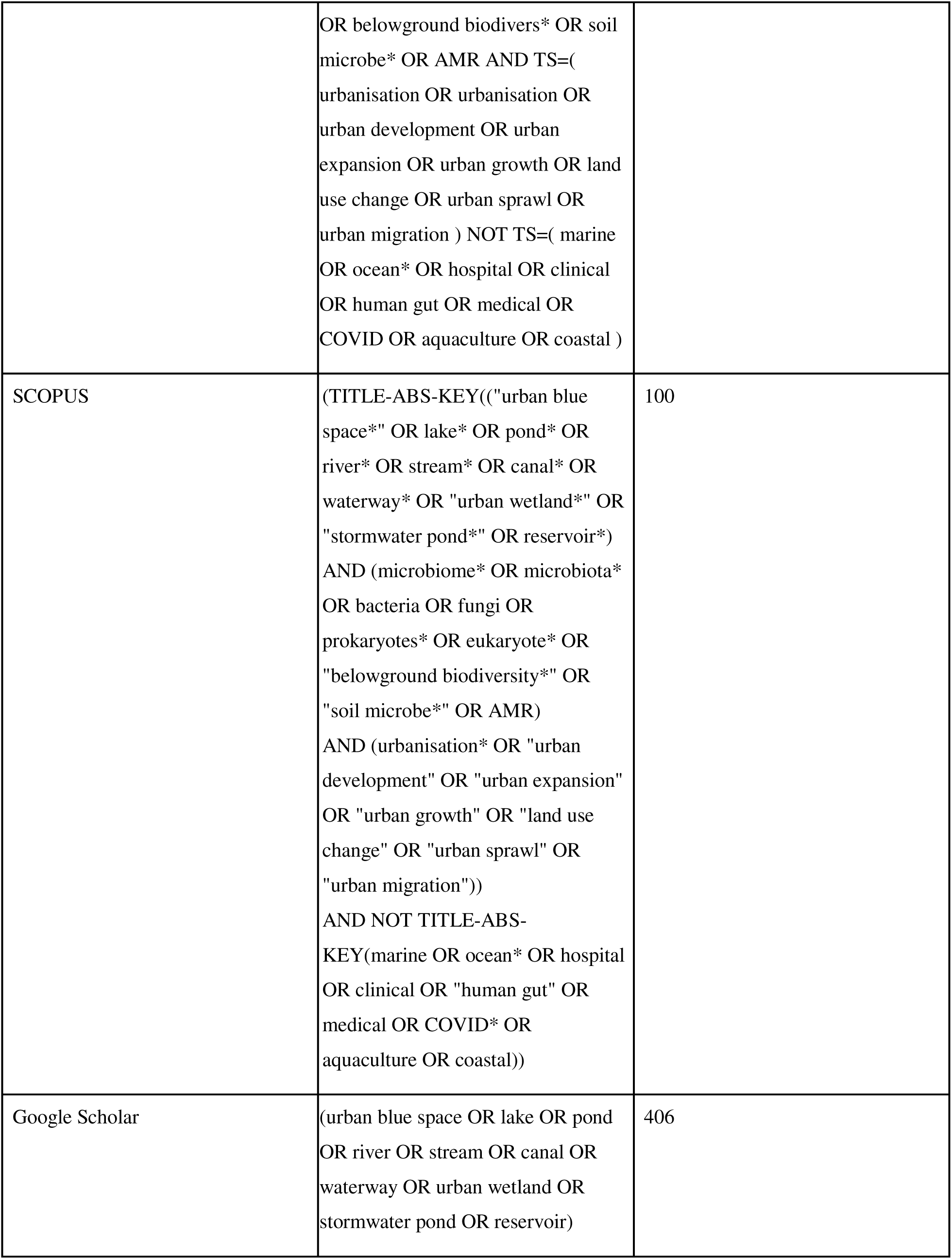
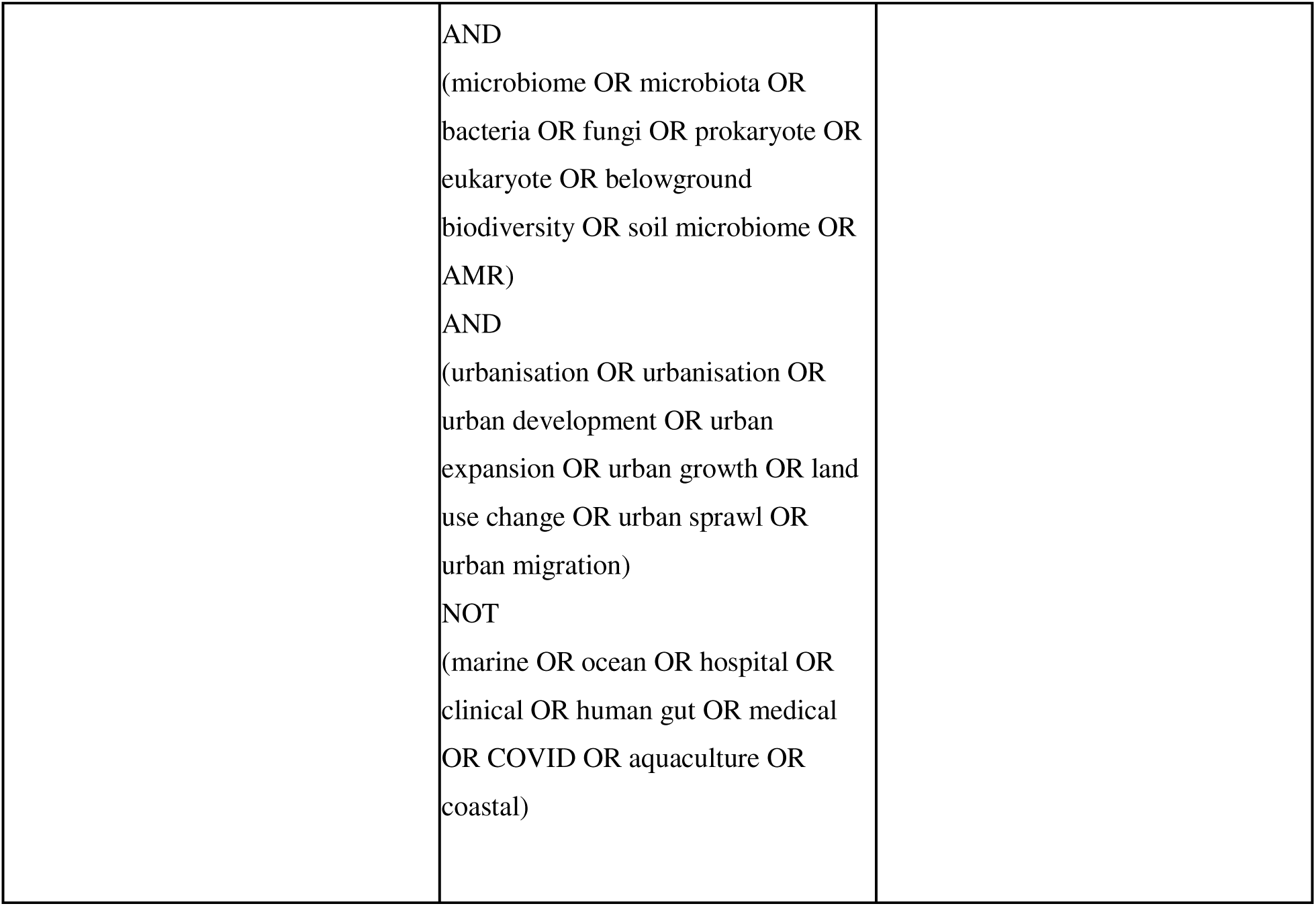
Search strings and number of records retrieved.

| Databases | Search Strings | Records retrieved |
| --- | --- | --- |
| WOS | TS=( urban blue space* OR lake* OR pond* OR river* OR stream* OR canal* OR waterway* OR urban wetland* OR stormwater pond* OR reservoir*) AND TS=(microbiome* OR microbiota* OR bacteria* OR fungi OR prokaryote* OR eukaryote* | 1055 |
|  | OR belowground biodivers* OR soil microbe* OR AMR AND TS=(urbanisation OR urbanisation OR urban development OR urban expansion OR urban growth OR land use change OR urban sprawl OR urban migration ) NOT TS=( marine OR ocean* OR hospital OR clinical OR human gut OR medical OR COVID OR aquaculture OR coastal ) |  |
| SCOPUS | (TITLE-ABS-KEY(("urban blue space*" OR lake* OR pond* OR river* OR stream* OR canal* OR waterway* OR "urban wetland*" OR "stormwater pond*" OR reservoir*) AND (microbiome* OR microbiota* OR bacteria OR fungi OR prokaryotes* OR eukaryote* OR "belowground biodiversity*" OR "soil microbe*" OR AMR) AND (urbanisation* OR "urban development" OR "urban expansion" OR "urban growth" OR "land use change" OR "urban sprawl" OR "urban migration")) AND NOT TITLE-ABS-KEY(marine OR ocean* OR hospital OR clinical OR "human gut" OR medical OR COVID* OR aquaculture OR coastal)) | 100 |
| Google Scholar | (urban blue space OR lake OR pond OR river OR stream OR canal OR waterway OR urban wetland OR stormwater pond OR reservoir) | 406 |
|  | AND<br>(microbiome OR microbiota OR<br>bacteria OR fungi OR prokaryote OR<br>eukaryote OR belowground<br>biodiversity OR soil microbiome OR<br>AMR)<br>AND<br>(urbanisation OR urbanisation OR<br>urban development OR urban<br>expansion OR urban growth OR land<br>use change OR urban sprawl OR<br>urban migration)<br>NOT<br>(marine OR ocean OR hospital OR<br>clinical OR human gut OR medical<br>OR COVID OR aquaculture OR<br>coastal) |  |

**Table 2:** Different inclusion and exclusion criteria that were used to select the studies meant for mapping. The criteria includes synonyms and the rationale behind each criteria.

| Inclusion | Rationale |
| --- | --- |
| Urban blue spaces | Studies conducted in freshwater bodies, lakes, rivers, ponds |
| Water body<br>microbe/microbiome/microbiota/bacteria/fungi/prokaryotes/eukaryotes | Studies involving microbes present in the water hence, can include bacteria, fungi, protists, some types algae, individual taxa and groups/community |
| Relevant ecosystem | Only aquatic freshwater |
| Analytical modelling | Correlative studies are included but analytical techniques used are also noted to understand the cause-effect relationship depicted |
| Time Period | 2000-2025 |
| Language | Studies published in English language or a reliable translation to English |
| Functional traits | Studies depicting functional traits such as antibiotic resistance, nitrogen cycling genes are also considered |
| Exclusion | Rationale |
| Urban blue spaces | Excluding spaces like coastal areas, bays, harbors, stormwater |
| Heavily industrialized areas, areas that have been left behind (like landfills and so on) | Excluding spaces that involved extensive industrialisation, because those sites will be far away from urban spaces and consist of heavy metals which may drastically alter the community. |
| Non-urban | Excluding studies involving non urban blue spaces present in rural areas or pristine areas. |
| Ecosystems | Excluding air-borne microbiomes, microbiomes on surfaces other than soil, endosphere, phyllosphere and wetlands. |
| Soil microbiome | Excluding studies involves the microfauna, meiofauna, macrofauna megafauna present in the soil. |
| Reviews/speculative studies/Human health | Excluding Article/literature reviews/meta-analysis and studies which focus on human health |

We used Google Scholar, Web of Science and SCOPUS to search for articles, develop the search string as well as the inclusion-exclusion criteria. The final search string used for each database can be found in Table 1. WOS, GS and SCOPUS were the databases used to obtain studies for analyses. In the search string, we used terms synonymous with lakes, waterbodies, urbanisation and microbiome. We also used Boolean operators such as “AND” and “OR”. We identified more articles by going through the bibliographies of the studies included after primary screening. For WOS and SCOPUS we retrieved all the results of each search string, for GS we retrieved results from the first 10 pages for each search string. These were collated and duplicates were removed.

The scoping exercise revealed that studies revolving around the water microbiome and urbanisation began around 2000 and so, we included research published from 2000 to 2025. We did not place any restriction on geographic locations but, we only considered published studies (i.e., no grey literature) and studies published in English as it was difficult to obtain reliable translations.

We estimated the comprehensiveness of the search string by checking for overlap between the three databases. We found a significant overlap between the three databases, suggesting that all three databases covered important studies published during the established time frame. The collated list of papers was divided into 5 groups such that every section was screened by two authors. This screening was based on the title and abstract. The agreement coefficient for each section was calculated. The agreement coefficient is a statistical measure used to quantify the reliability between two raters using different methods. In this study, the method used is Cohen’s kappa. The kappa value was calculated for each section and overall value (0.85) was obtained by averaging those values. This was followed by full-text screening.

The data extracted (Table 3) was: microbial communities, microbial indices, microbial functional traits, sample information, site information, study site details, climatic conditions, techniques used for microbial identification, type of urban water body, type of human activity observed or studied, relationship between urbanisation and microbes, types of statistical tests, electronic publishing date, journal the study is published in, authors origins and funding agencies involved. The results were through graphs and the graphs were made using R and QGIS.

**Table 3:**
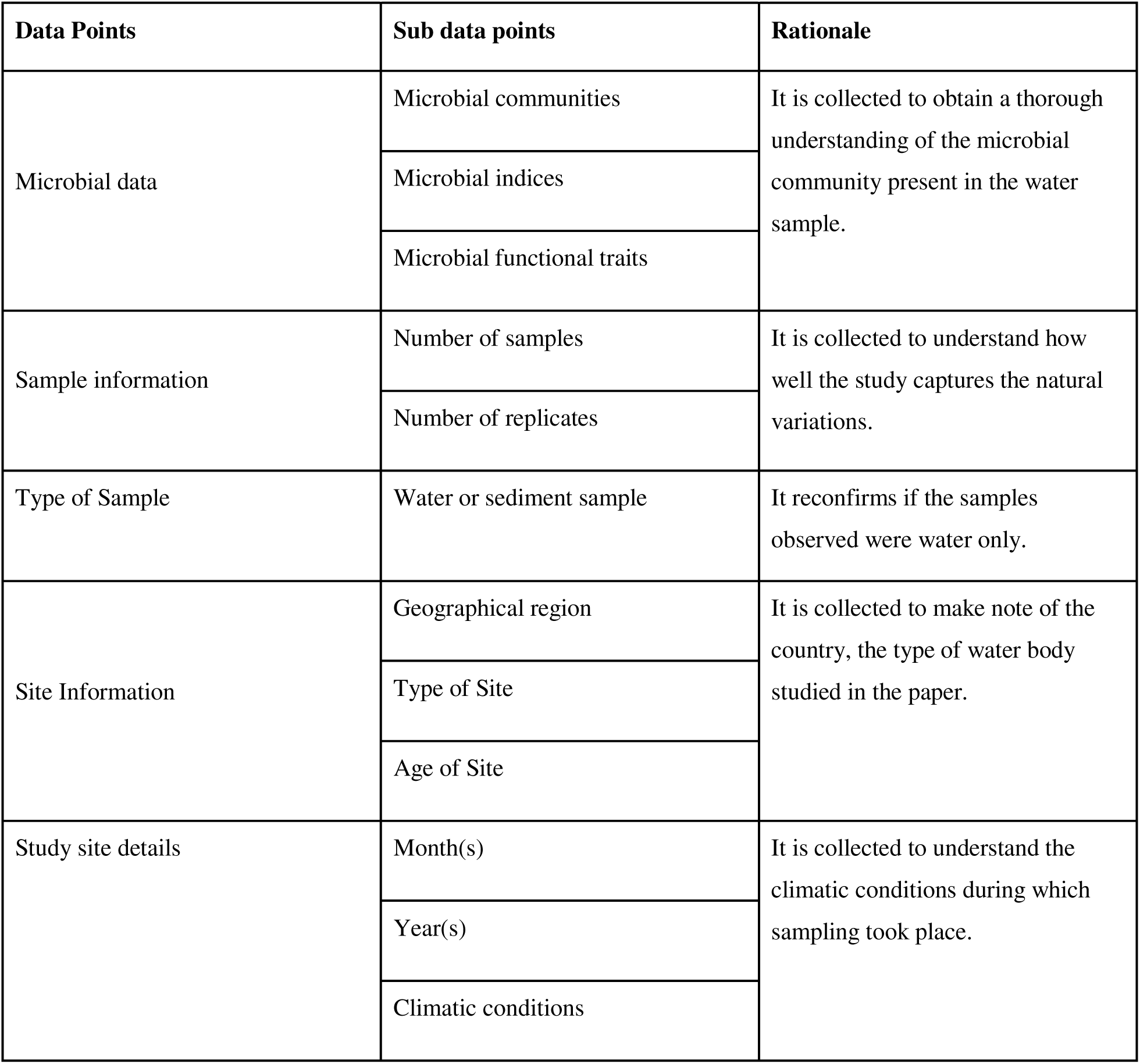
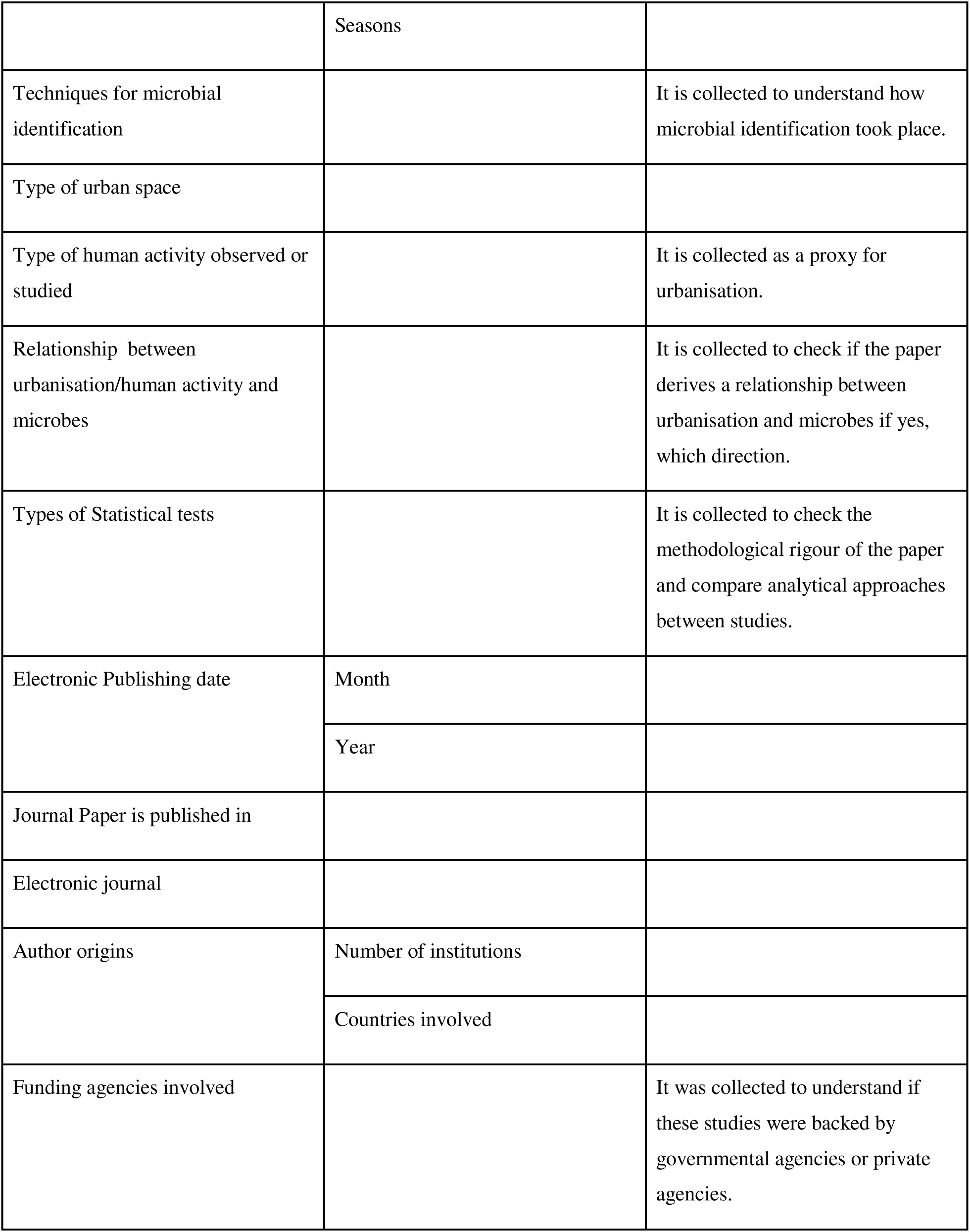
Each data point that was extracted along with subcategories for each. It takes into account the variability observed while scoping.

## Result

### Number of Studies

The search string resulted in 1,561 papers from three unique databases. After removing duplicates, 1,059 papers were screened. 940 papers were excluded after title and abstract screening from the review based on the eligibility criteria. Out of 123 papers, 17 papers could not be retrieved. 106 papers were screened for full text screening, out of which 16 rejected further based on the exclusion criteria. Thus, 90 papers were included in the review. Fig 1 shows the PRISMA (Page et al., 2020) chart for details of the screening process.

**Fig. 1.** PRISMA flow chart for systematic literature reviews. The flow chart shows the systematic approach taken to select the studies for the review. It shows the number of studies selected, removed and reason of exclusion at each step. The agreement coefficient was calculated after the primary screening (Page et al., 2012).

### Geographic Distribution of Studies

There is an uneven geographic distribution of papers across the globe depicted in Fig 2. Developed countries such as China, United States of America and Canada are over represented in water microbial research. China contributed the highest number of studies (55.5%, n = 50) followed by the United States of America (12.22%, n = 11, Fig 3). South Asia and Africa had fewer studies on urban freshwater microbiomes, despite facing more water stress due to rapid urbanisation and industrialization.

**Fig 2.**
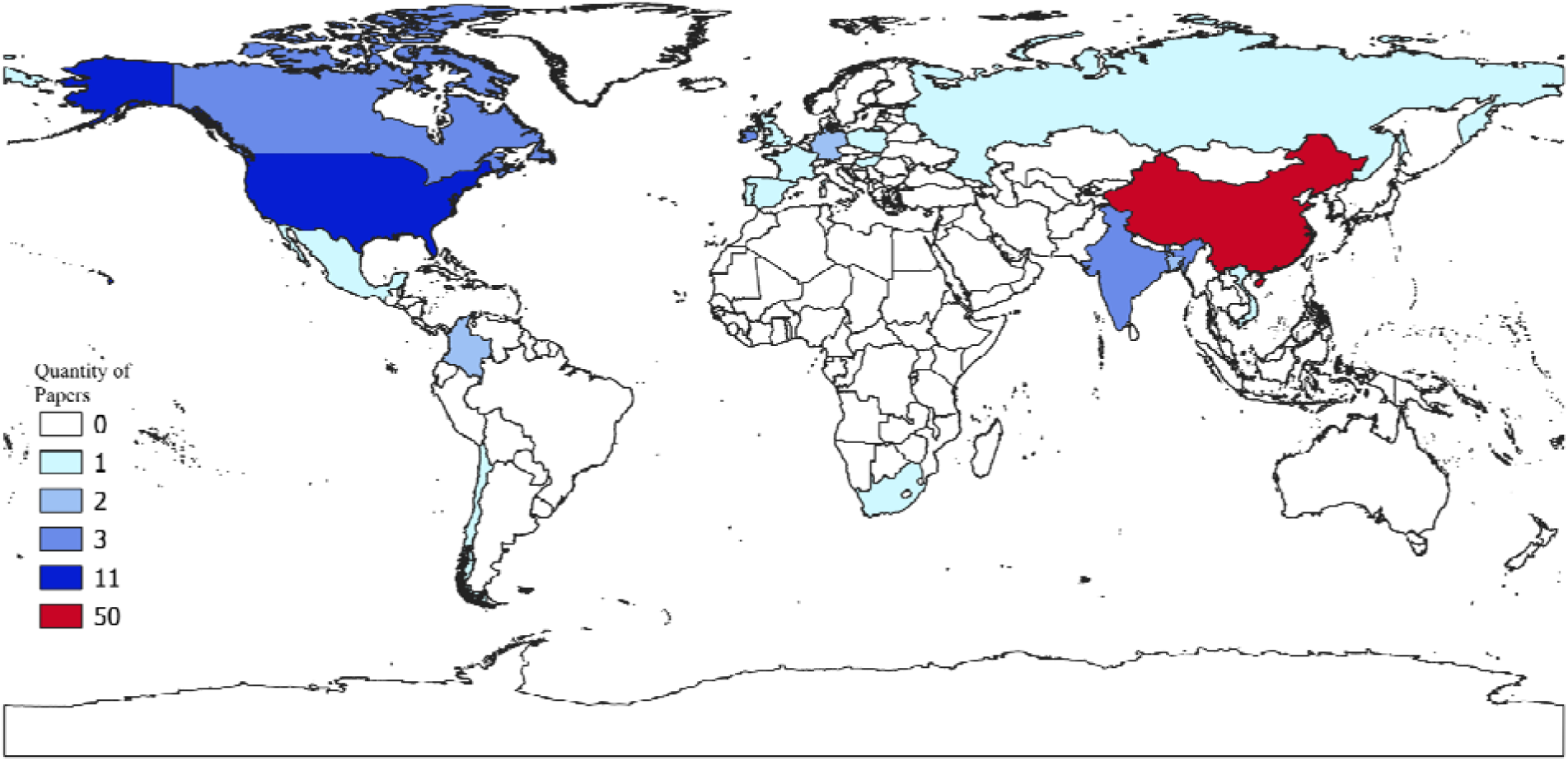
The global distribution of studies on urban freshwater microbiomes. More than 50 studies have been conducted in the People’s Republic of China followed by the USA with 11 studies.

**Fig 3:**
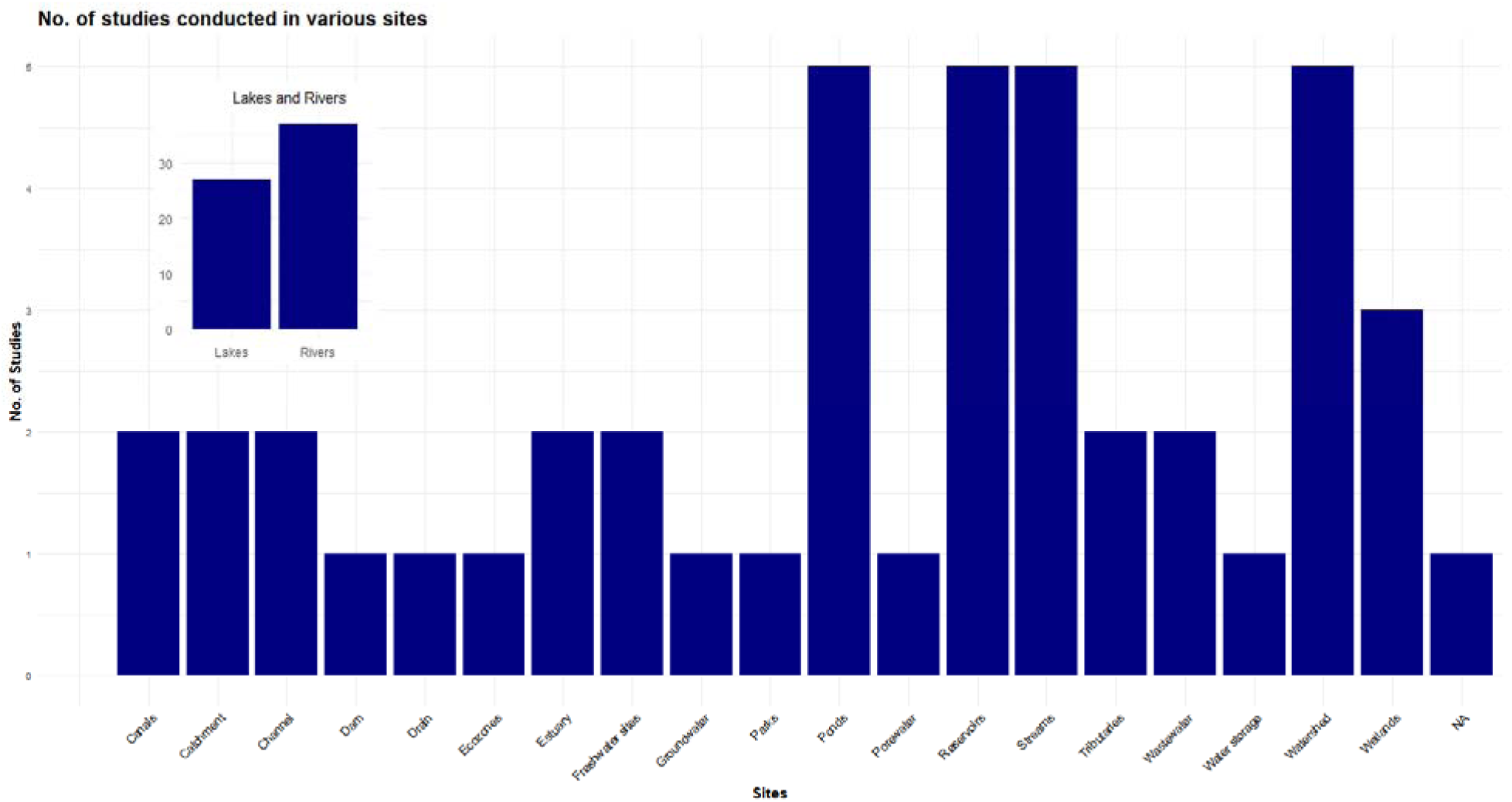
This figure shows the number of studies conducted in different types of sites. Freshwater sites also include Freshwater springs. NA in the graph, is for the studies that did not mention the type of sites just written as water samples taken from surface waters. Waste water includes influents and effluents of the wastewater system. And the Rivers also include the riverine system.

### Study Sites

Among the selected papers, lakes and rivers are the most commonly studied sites (n=27 and 37) respectively, showing proportionally higher attention. In contrast, the other types of sites such as channels, estuary, catchment areas as depicted in Fig. 3 are studied much less frequently, appearing in only one or two papers. Few studies among the examined publications include water samples from nearby construction sites that show the direct impact of anthropogenic activities on waterbodies and the microbiome inside it.

### Microbial Functional Traits Studies

Nutrient cycling was the most frequently reported functional trait, appearing in 30.8% of included studies(Fig. 4), contributing 30.80%, followed by AMR/pathogen-related traits at 20.09%. Stress-related traits, such as seasonal variation, biofilm formation, and temperature tolerance, accounted for 14.73%. In contrast, physical traits and mobile genetic elements/horizontal gene transfer were underrepresented.

**Fig 4.**
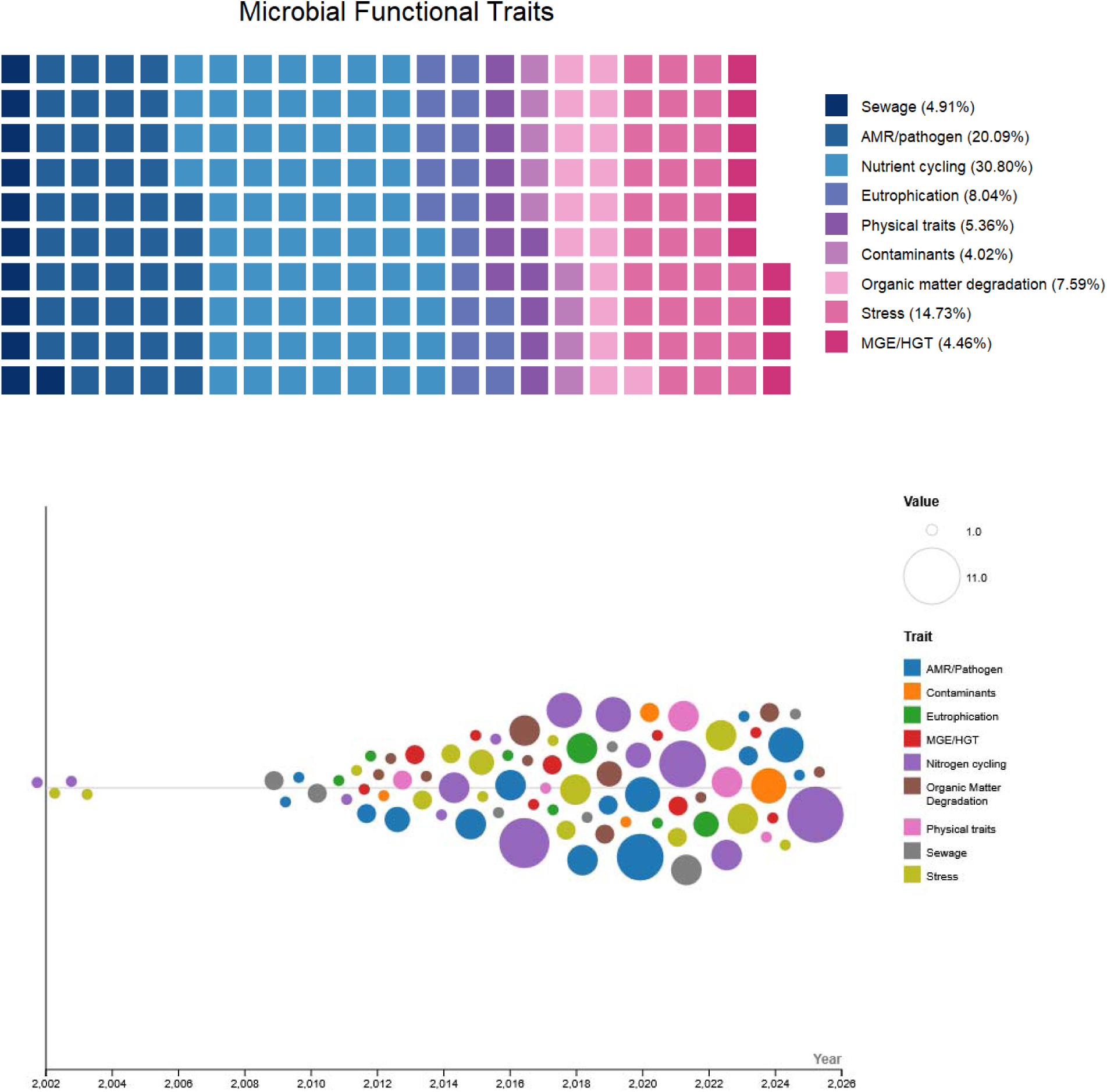
**a.** The above figure shows the frequency of various microbial functional traits occurring among the selected papers. **b.** The frequency of various functional traits being studied each year.

The temporal look at the functional trait, Fig 4b, shows that studies on microbiome were limited until 2010 but have grown exponentially since 2016, and all traits have been studied relatively evenly across the years since.

### Anthropogenic Activities

The most studied anthropogenic activity across the studies were tourism and tourist residence (21.6%, n = 106, Fig 5), followed by land use impact (20%, n = 100), sewage (16.2%, n = 81) and discharge (12.8%, n = 64) as shown in Fig 5. Because studies often report multiple drivers, percentages represent frequency of mention rather than mutually exclusive categories.Certain drivers such as stormwater runoff and cultural practices appear underrepresented, suggesting potential gaps in current research focus.

**Fig 5.**
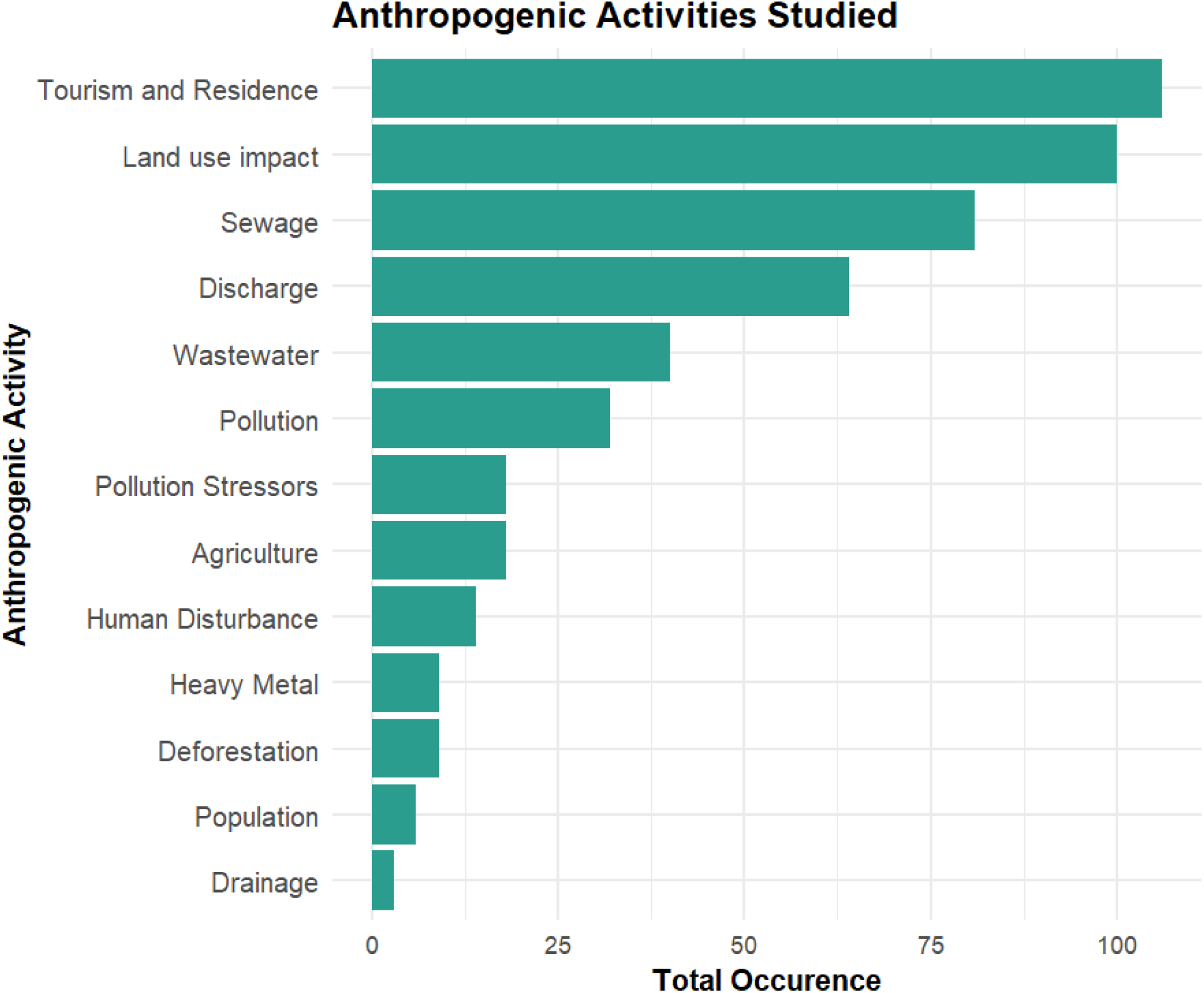
The above figure shows the frequency of various anthropogenic activities studied in the selected papers. Tourism and residential land use were the most frequently studied anthropogenic drivers, followed by land-use change, sewage discharge, and wastewater inputs.

### Microbial Diversity

Many studies reported the presence of bacteria without further classification. The most commonly reported bacterial groups were Proteobacteria, followed by Cyanobacteria and Actinobacteria(Fig 6). Other frequently reported groups included Firmicutes, Chloroflexi, and Bacteroidetes. Some of the least reported groups were Deferribacteres, Frankiales, Parcubacteria, and Thaumarchaeota, as depicted in Fig. 6.

**Fig 6.**
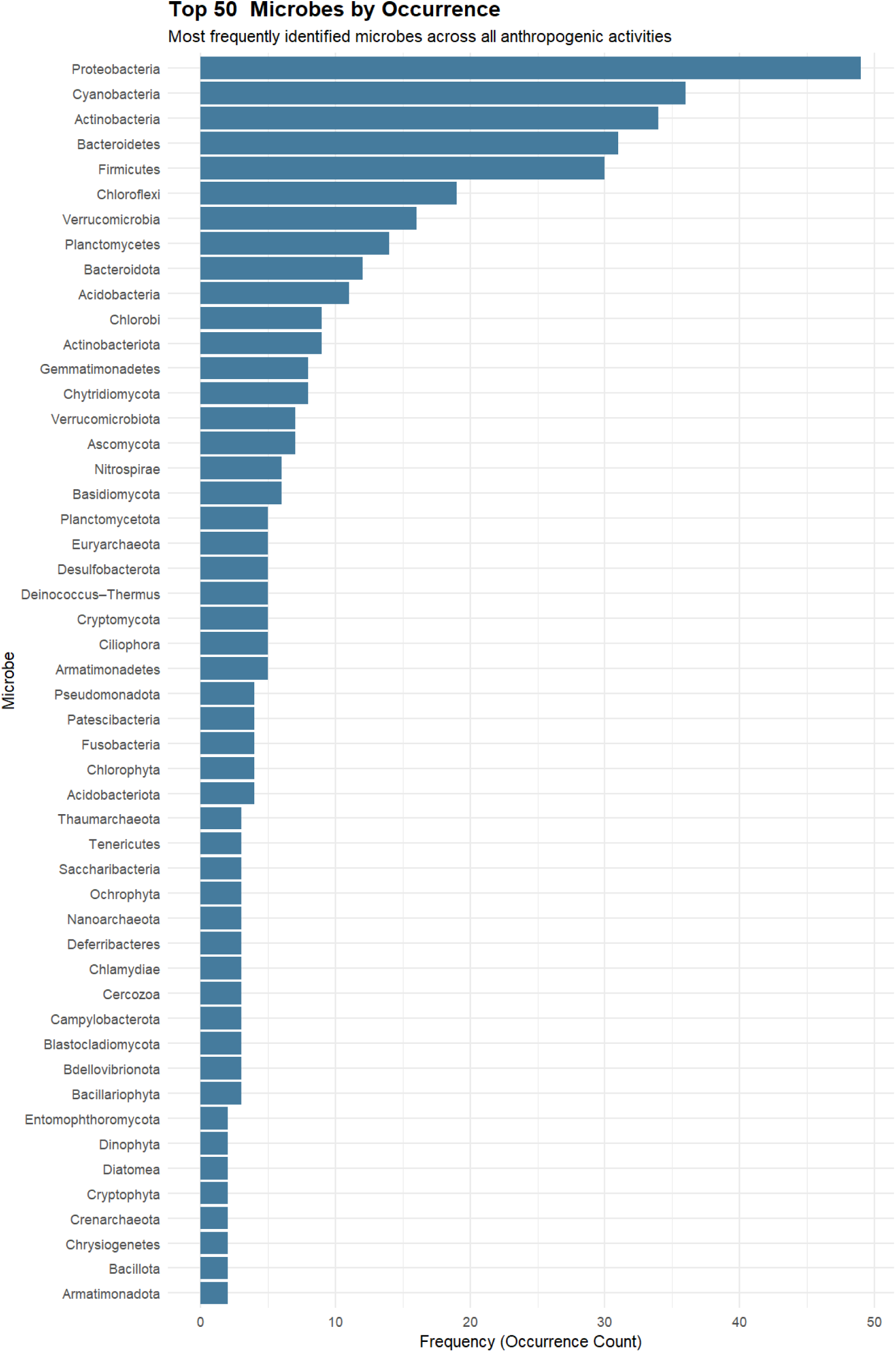
The top 50 microbes classes found in the papers. Proteobacteria is a broad generalist bacterial phylum that is present in almost all the papers studied. It was followed by Cyanobacteria, Actinobacteria, Bacteriodetes, Firmicutes and Chloroflexi.

It represents the frequency of reporting across studies and does not necessarily reflect relative abundance changes under urbanisation. The most frequently reported microbes were microbes from the human gut, indicating a sewage contamination in the waterbodies. Further, many of the reported microbes have proven to have harmful impacts on human health, such as Proteobacteria, having evidence in metabolic disorders and inflammatory bowel disease and playing a role also in lung diseases, such as asthma and chronic obstructive pulmonary disease (Rizzatti et al, 2017). Cyanobacterial blooms result in the bioaccumulation of cyanotoxins that are associated with renal failure, decreased platelets, increased leukocytes, and the development of hepatotoxicosis (Villalobos et al, 2025).

### Microbial Diversity across Anthropogenic Activities

The heatmap reflects co-reporting patterns of microbial taxa across studies examining different anthropogenic drivers. Fig 7 does not represent quantitative abundance measurements within individual waterbodies. Tourism, Discharge, Pollution, Wastewater, Land Use, and Sewage)shows significantly higher microbial abundance and diversity. Proteobacteria and Actinobacteria, generalists, show positive association with sewage, tourism and land use impact, suggesting they play an important role in the formation of the particular microbial signature of a water body. Agriculture, Deforestation, and Drainage promote the establishment of a few more niche or specialist species due to the conditions provided. Activities like Sewage, Land Use, and Pollution are clustered together because they all show high microbial abundance. On the other hand, Agriculture and Deforestation are branched off alone because their microbial communities are different. This could be due to the uneven distribution of papers across all anthropogenic activities, but this still highlights how different anthropogenic stressors mold the microbial signatures of various ecosystems.

**Fig 7.**
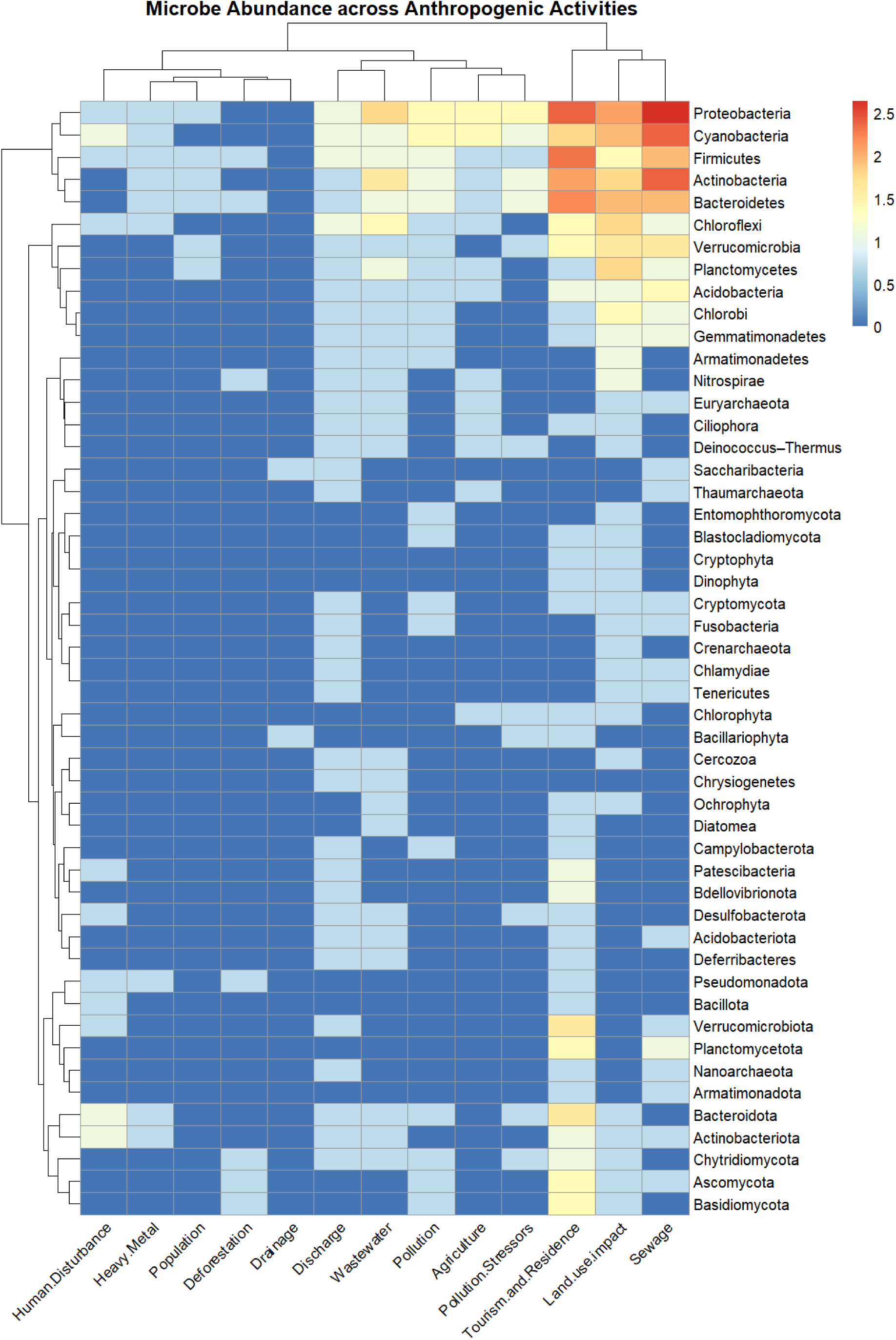
The above heat map shows the presence of microbial communities between various anthropogenic activities. The bacterial count is taken in logarithmic scale to show the variation between the count. Generalist taxa such as Proteobacteria, Cyanobacteria, and Actinobacteria were consistently reported across studies examining multiple anthropogenic activities. The groupings show the anthropogenic activities as well as microbial community composition that are the most identical to one another.

### Effect of urbanisation across studies on microbial diversity

Urbanisation was consistently found to be a driver of microbial community restructuring, and this was evident both taxonomically and functionally (Table 4 and 5). Alpha diversity (28 studies) was found to be inconsistent and therefore not a reliable measure of urbanisation. Diversity was found to be maintained or even enhanced under nutrient enrichment, even where changes to the microbial community were evident (26 studies). These changes were found to be more indicative of the effects of urbanisation. Proteobacteria was dominant (33 studies), and this was enhanced, especially in wastewater and nutrient-rich environments. This was due to the ability of this group to thrive under disturbed conditions. Conversely, coliforms and E. coli (18 studies) were found to increase significantly in urbanized environments, especially where there was a sewage input. This was a reliable indicator of urbanisation. Pathogens and virulence factors (11 studies) were also found to be enriched.The functional response was found to be closely associated with the urban drivers. The antibiotic resistance genes (ARGs) were found to be widespread and enriched, especially in wastewater-influenced systems, suggesting that urban waters harbor antimicrobial resistance. The nitrogen cycling genes and associated microorganisms were found to be consistently altered, suggesting the enhanced nutrient transformation potential under eutrophic conditions. Similarly, the carbon metabolism was found to be enriched in response to organic pollution and DOM, suggesting enhanced microbial carbon transformation potential. Overall, the functional potential was found to be enriched in response to urban drivers, including pathways associated with respiration, nutrient cycling, and stress response.

**Table 4.**
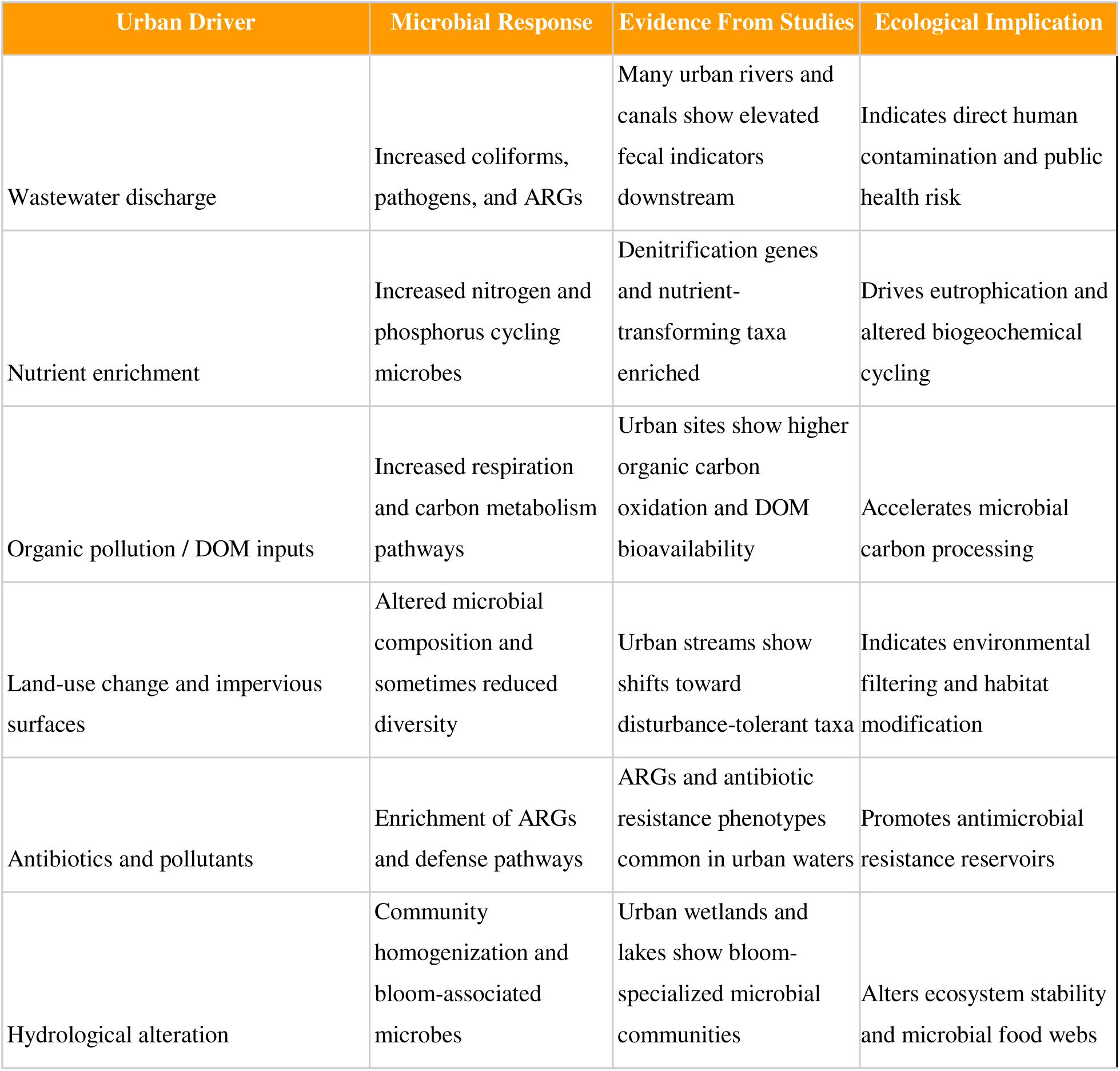
Different types of urban drivers explored across the 90 studies that alter microbial communities, increasing pathogens, ARGs, and nutrient-cycling taxa, with implications for ecosystem function and public health.

| Urban Driver | Microbial Response | Evidence From Studies | Ecological Implication |
| --- | --- | --- | --- |
| Wastewater discharge | Increased coliforms, pathogens, and ARGs | Many urban rivers and canals show elevated fecal indicators downstream | Indicates direct human contamination and public health risk |
| Nutrient enrichment | Increased nitrogen and phosphorus cycling microbes | Denitrification genes and nutrient-transforming taxa enriched | Drives eutrophication and altered biogeochemical cycling |
| Organic pollution / DOM inputs | Increased respiration and carbon metabolism pathways | Urban sites show higher organic carbon oxidation and DOM bioavailability | Accelerates microbial carbon processing |
| Land-use change and impervious surfaces | Altered microbial composition and sometimes reduced diversity | Urban streams show shifts toward disturbance-tolerant taxa | Indicates environmental filtering and habitat modification |
| Antibiotics and pollutants | Enrichment of ARGs and defense pathways | ARGs and antibiotic resistance phenotypes common in urban waters | Promotes antimicrobial resistance reservoirs |
| Hydrological alteration | Community homogenization and bloom-associated microbes | Urban wetlands and lakes show bloom-specialized microbial communities | Alters ecosystem stability and microbial food webs |

### Correlation of Microbial Diversity across anthropogenic activities

Fig 8 highlights how different anthropogenic stressors mold the microbial signatures of various ecosystems. The correlation matrix shows the statistical relationship between the microbiome of various anthropogenic stressors. Spearman’s Correlation was used to calculate the correlation coefficients.

**Fig 8.**
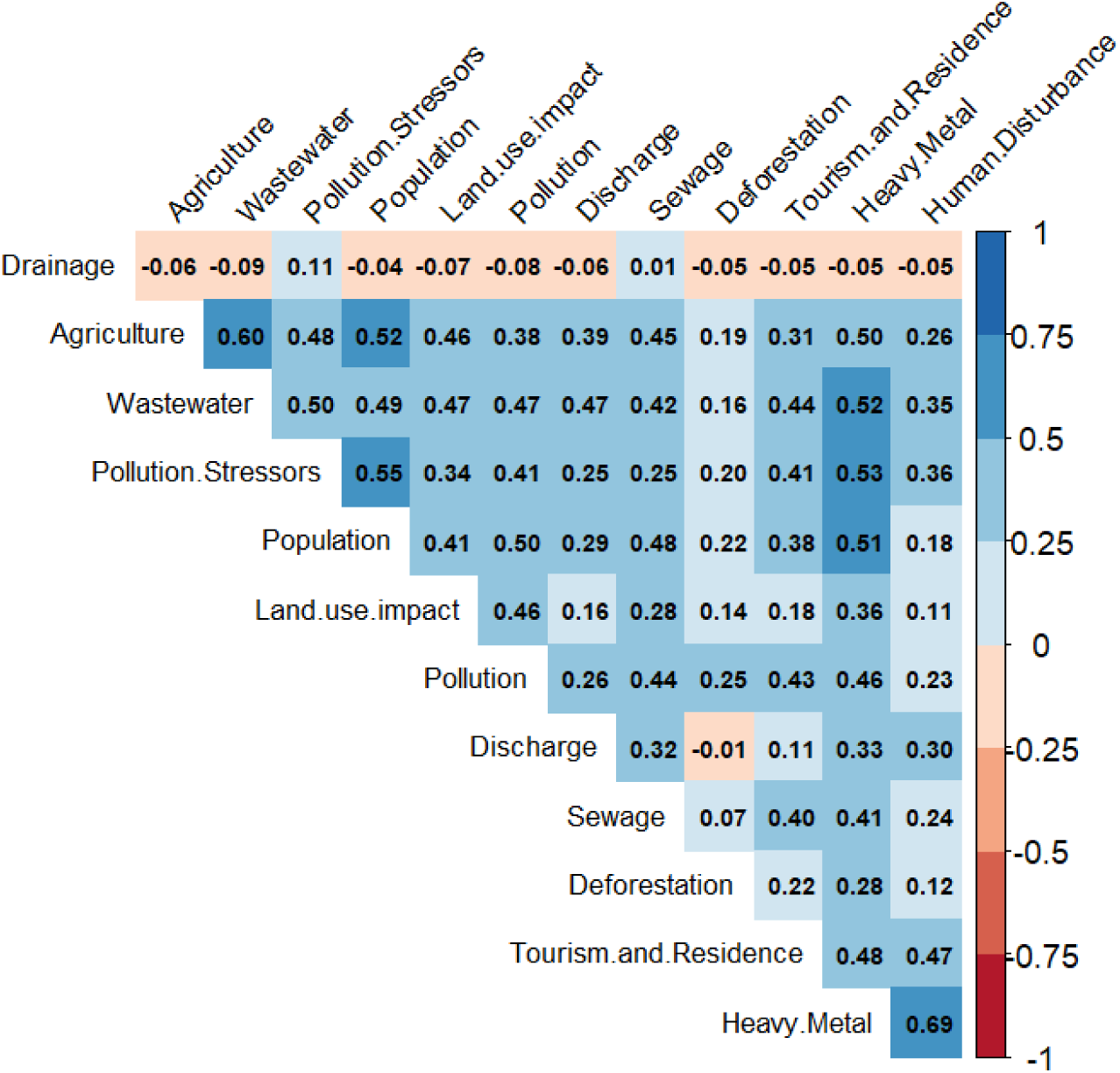
The correlation plot between the microbial community found across various anthropogenic activities. The Spearman’s coefficients are given between the microbial communities most reported in presence of anthropogenic activity. The strongest correlation was found in agriculture with wastewater and population, heavy metal with human disturbance, wastewater, pollution stressor and population. Higher positive correlation indicates presence of similar microbial communities in both the anthropogenic activity sites. The correlation coefficients between all the activities show higher positive correlation due to presence of generalist species such as Proteobacteria. This reflects similarity in taxa reported across studies rather than direct ecological interactions within individual waterbodies.

The strongest positive correlation was found between Heavy Metal and Human Disturbance. This suggests that patterns of human distribution are strongly correlated with levels of heavy metal contamination in urban waterbodies, and are associated with shifts in the native microbiome. Agriculture and wastewater with its high positive correlation indicate that the farming structure shares the same basin for wastewater discharge and wastewater. Drainage has the most independent microbiome composition, not dependent on the other anthropogenic activities listed. The correlation matrix has more positive correlations due to the higher abundance of generalists such as Proteobacteria and Actinobacteria, which drives homogeneity across sites. The correlation analysis reflects similarities in reported microbial taxa across studies investigating different anthropogenic drivers and should not be interpreted as within-site ecological interactions.

### Principal Component Analysis

In Fig 9 Dim1 represents the “total human impact or gradient of urban pollution”. Samples on the far right are highly contaminated; samples on the left are relatively “clean”. Dim1 has a high explanatory power of 55.5% tells us that microbial diversity across waterbodies is driven by the Human Impact gradient. Dim2 represents the secondary variation distinguishing between urban and population-related impacts and industrial impacts. Sewage, Pollution, Wastewater, and Tourism/Residence are packaged together and directed towards the right, which indicates that they are correlated and drive similar microbial changes in waterbodies. Heavy Metal and Human Disturbance point downward, suggesting they create a distinct environmental niche that differs from “Sewage”. Drainage and Deforestation point in opposing directions from the main cluster indication; they have a unique effect on the aquatic environment that is not completely exhaustive with the urban pollution. The longer the arrow represents more importance of the stressors in shaping the microbial communities of the waterbodies. Proteobacteria, Cyanobacteria, and Actinobacteria are highly “enriched” by urban waste. They likely thrive in high-nutrient (eutrophic) conditions provided by sewage and tourism-related runoff. The microbes in the central area are not strongly associated with these specific stressors.

**Fig 9.**
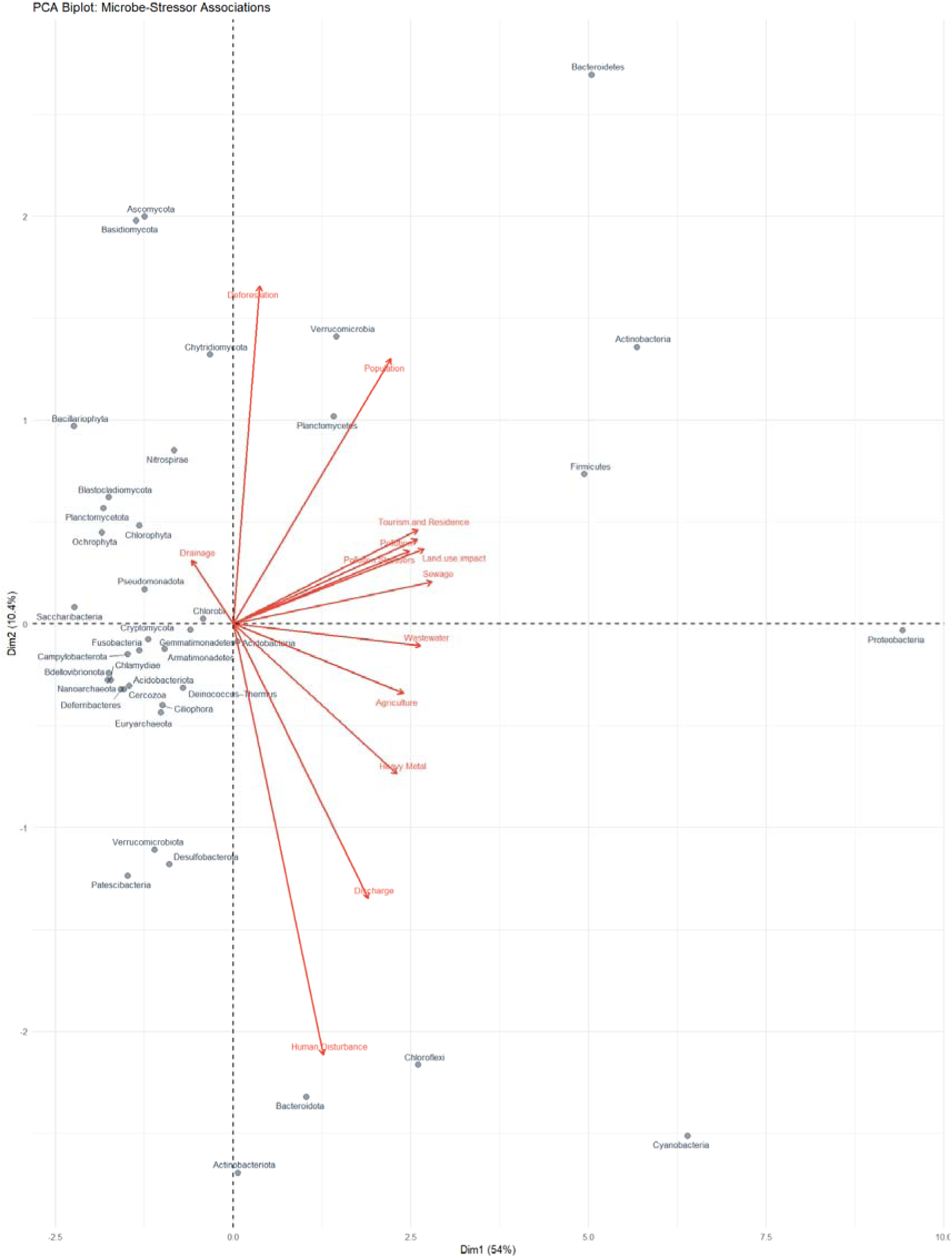
PCA analysis revealed clustering of anthropogenic drivers such as sewage discharge, land-use change, and pollution stressors, indicating similarity in microbial taxa reported across these disturbance categories.PC1 = disturbance intensity, PC2 = functional variation.

### Relative Stressor Composition

The Relative Stressor Composition, Fig 10, reveals that the most dominant taxa are heavily partitioned by anthropogenic pressures. Proteobacteria, Cyanobacteria, and Actinobacteria show larger sections for sewage, tourism and wastewater. High percentages of these blocks suggest these microbes are robust indicators of urban runoff. This shows the microbial community composition in urban waterbodies is not random but is specifically composed by the type of environmental stressors present.

**Fig 10.**
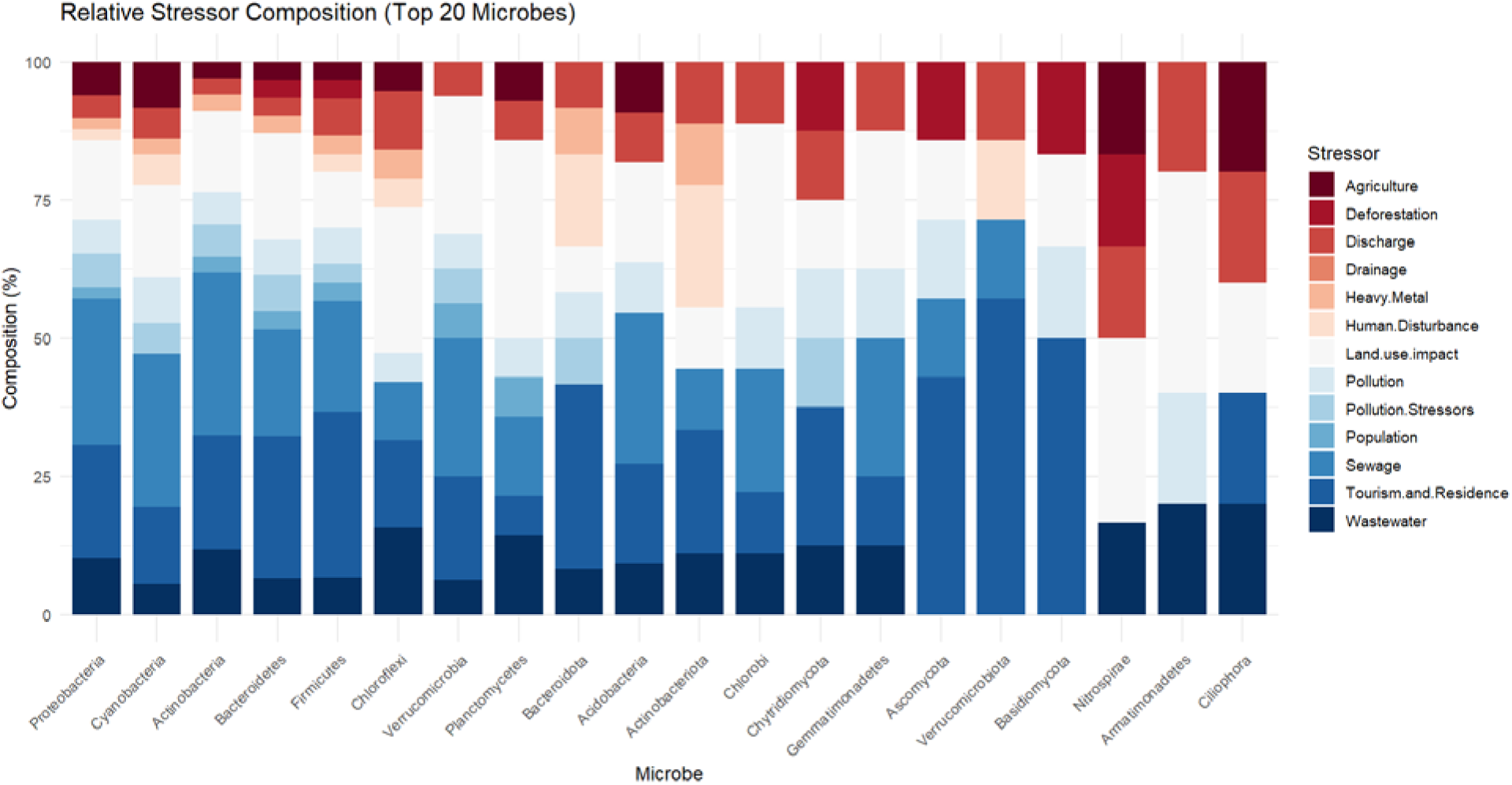
Relative Stressor composition of top 20 microbes. Proteobacteria, Cyanobacteria and Actinobacteria are comprised of multiple stressors with various relative weightage. But some like Ciliophora, Armatimonadetes and Nitrospirae have fewer stressors indicating they are present only in certain ecosystem conditions.

### Limitations

One major limitation of this study is the lack of rural urban comparisons in the studies that we picked up based on our search strings, which would have made the study stronger and more comprehensive. We were also forced to rely on reporting frequency of different microbes which may have resulted in over dominance of certain species across studies; while useful for detecting major trends, this method does not take into account effect size, magnitude of change, or significance, and therefore certain taxa may be emphasized while others, though showing small changes, may not receive adequate representation. With the timeline we selected for this review there is considerable heterogeneity between sequencing platforms, depth of sequencing, and bioinformatic pipelines, such as different regions of 16S rRNA, and these differences make studies less comparable to each other, which could lead to biases in community composition, diversity, and taxonomic resolution. Additionally, with most of the studies reported from China and North America our data gets skewed due to limited representation from the Global South, and this bias constrains the generalizability of findings, particularly given that urbanisation patterns, infrastructure, and environmental conditions differ substantially across regions. Finally, we chose papers which were written in English, which may have resulted in language bias, as studies conducted in other languages, especially in countries experiencing rapid urbanisation, may have been excluded, leading to an incomplete global overview due to the lack of local studies.

## Discussion

This is a comprehensive systematic review of the current global literature available on the influence of urbanisation on microbial communities in freshwater bodies. From 90 selected papers we identified critical gaps and biases that hinder a holistic understanding of these interactions. Notably, there is a lack of research from economically developing regions, Africa, and Southeast Asia (Fig. 2), limiting our ability to generalise global findings or look for geographical patterns in these areas. This is particularly crucial given that the predicted AMR burden in these regions is expected to be highest by 2050 especially affecting Africa and South Asian countries(Naghavi et al., 2024). In this context, the functional and taxonomic patterns identified in previous research become a key factor in understanding the ecological aspects of this risk.

A high prevalence of functional traits related to nutrient cycling and AMR/pathogens were identified, the co-occurrence of which suggests that urban freshwater systems are shaped by resource enrichment and may serve as petri dishes for resistance proliferation behaviour. The reporting frequency of Proteobacteria,Cyanobacteria, Actinobacteria, and Firmicutes adapt well to nutrient-rich and organically loaded environments. This also mirrors well to what research studies have reported over the last 25 years. Additionally, these phyla are ubiquitous in areas of high anthropogenic pressure where freshwater are often at the receiving end of sewage, untreated wastewater, runoff, and pollutants thus selecting for generalist microbial groups capable of rapid growth and metabolic flexibility. This is consistent with studies from Germany that show (Daga Quisbert et al., 2021) how urbanisation poses long-term health risks in waterbodies.

Similarly, in Numberger et al. (2022), Actinobacteriota and Gammaproteobacteria had ASVs unique to urban waters in freshwater bodies, including *Acidovorax* (Burkholderiales), *Flavobacterium* (Bacteroidota), and *Pseudomonas* (Gammaproteobacteria). Mohanta and Goel (2014) found that multiple drug-resistant bacteria were highest in samples from rapidly urbanising areas. Urbanisation thus acts as an ecological filter, with similar microbial patterns emerging globally in climatically distinct regions, selecting for phyla that rapidly grow in areas affected by sewage and tolerant of disturbances such as pollution thereby favouring opportunistic pathogens.

Above mentioned biases aside, due to the limited number of studies from the Global South, the methodology of different studies is also varied. Some studies have relied on single-season, short-term sampling or screening, as compared to systematic sampling over different seasons (Wang *et al* 2018, Zhao *et al*. 2022). It is difficult to infer the effect of seasonal variations on microbiome diversity. Certain studies have also derived their data from 16s amplicon sequencing, while few others from metagenomic analysis. There is a significant gap in the availability of transcriptomic, proteomic, and metabolomic data. Although some research has examined samples across urban gradients, most studies are restricted to a limited number of sampling sites, making broader ecological inferences challenging.

The consistent reporting of similar bacterial phyla across diverse geographic regions and anthropogenic disturbances suggests increasing microbial homogenisation in urban freshwater ecosystems globally.China, which has the most number of studies on urban water microbiomes, is suffering from waterbody loss (Xiao et al., 2022). This is mainly due to urbanisation, where waterbodies play an important role in providing drinking water. China being one of the world’s leading producers and consumers for antibiotics, is a potential centre for development and dissemination of ARGs (Zhang et al., 2023). As a precautionary measure, they have invested a lot of funding towards harmonious development between resources and environments, in the form of ‘Sponge city programs’ and ‘The Administrative Measures of the Urban Blue Line’(Xiao et al., 2022).

Similarly studies on freshwater microbiomes need to be expanded to the economically developing countries as these studies reveal the presence of human health risks, impacts of pollution and help in the long term monitoring of the ecosystem health (McLellan et al., 2015). The global understanding of urban freshwater microbiomes is currently geographically skewed, limiting predictive power and policy relevance in regions expected to face the highest AMR burden by 2050.

As a finite resource it is vital to maintain the microbial biodiversity of urban waterbodies to sustain people. This also brings up questions around equity as marginalised communities depend more on freshwater bodies for survival. In a (Satterthwaite et al. 2022) Bengaluru case study, the most rapidly urbanising city in India, it was discovered that lower income communities closer to the lake faced higher risk of water related fecal contamination. Community distribution of microbes, if conducted in countries of South America and South Asia, can help us estimate the disease vulnerability of communities that depend on such wetlands for survival. Public health welfare programs and education campaigns around the microbial make-up of an ecosystem like waterbodies especially with AMR on the rise can create some much needed awareness around these sites.

As Maria Magdalena Warter, points out “Just like our gut, freshwater ecosystems need a functioning microbiome. Bacteria and other microorganisms form the basis of food chains and metabolic processes, as well as the self-purification capacity of waterbodies”(Leibniz Institute of Freshwater Ecology and Inland Fisheries (IGB), 2025). Thus there is certainly a need for a better wastewater management system that protects the natural urban waterbodies and prevents further entry of AMR. Across Table 4 and 5 the findings were consistent with the expected effects of specific urban drivers. Antibiotics and pollutants are associated with enriched resistance traits, and hydrological changes were associated with homogenization and bloom-forming microorganisms.

**Table 5.** Urbanisation alters microbial diversity, composition, and function, with consistent increases in fecal indicators, ARGs, and disturbance-tolerant taxa, while community composition and functional potential shift more reliably than diversity alone.

| Indicator | Direction of Change | Number of Studies Reporting | Key Observations Across Studies | Ecological Interpretation |
| --- | --- | --- | --- | --- |
| Alpha diversity (Shannon, Chao1, richness) | Mixed | 28 | Some urban waters show lower diversity due to stress and pollution, while | Diversity alone is not a consistent indicator of urban impact; |
|  |  |  | nutrient enrichment and runoff can increase diversity. Many studies show stable diversity but major compositional shifts. | community composition is more sensitive. |
| Proteobacteria abundance | Frequently dominant | 33 | Proteobacteria commonly remain dominant across sites and often increase in disturbed waters, wastewater-influenced systems, and nutrient-rich environments. | Highly adaptable taxa associated with organic pollution, nutrient enrichment, and disturbance tolerance. |
| Antibiotic resistance genes (ARGs) | Increased or detected frequently | 19 | ARGs and resistance phenotypes detected in sediments, water, and biofilms; some studies show strong ARG–mobile genetic element correlations. | Urban waters may act as reservoirs and transmission pathways for antimicrobial resistance. |
| Coliforms / *E. coli* / fecal indicators | Strong increase in urban sites | 18 | Highest levels downstream of sewage outflows, wastewater discharge, and highly urbanized catchments. | Reliable indicators of anthropogenic contamination and sewage influence. |
| Pathogens / virulence factors | Often enriched | 11 | Opportunistic pathogens and virulence genes frequently detected in urban rivers, ponds, and canals. | Urban aquatic systems may act as pathogen reservoirs with potential public health implications. |
| Nitrogen cycling genes | Altered (often more denitrification or N-processing) | 15 | Changes in genes such as *narG*, *nirS*, *amoA*, and enrichment of nitrogen-transforming bacteria observed along urban gradients. | Nutrient enrichment and runoff alter biogeochemical cycling and ecosystem functioning. |
| Carbon metabolism / DOM processing | Altered | 13 | urbanisation changes DOM composition, respiration rates, and microbial carbon metabolism pathways. | Urban runoff and organic inputs reshape carbon processing and microbial energy pathways. |
| Community composition | Strong shifts or homogenization | 26 | Urban gradients drive major restructuring of microbial assemblages, even where alpha diversity remains stable. | Environmental filtering and pollution select for disturbance-tolerant microbial taxa. |
| Functional potential / metabolic pathways | Frequently altered | 17 | Enrichment of metabolic pathways related to respiration, nutrient processing, and stress response. | Indicates functional reorganization of microbial ecosystems under urban pressure. |

Urban freshwater systems increasingly reflect wastewater-associated microbial signatures, indicating that anthropogenic inputs contribute to a partial convergence between natural and human-associated microbiomes(Numberger et al., 2022). One can infer from this study that ARGs and fecal indicators may serve as more reliable monitoring targets than diversity metrics alone. These patterns reinforce a One Health perspective, where environmental microbial dynamics, ecosystem functioning, and human health risks are closely interconnected through shared exposure pathways. Policies around better wastewater and runoff management along with pathogen indicators can help mitigate AMR and associated risks. Future research should focus on using expansive research methods and a multi-omics approach, aside from developing standard methodologies and geographic representation, especially in rapidly urbanising areas. These research methods are going to be crucial in developing futuristic frameworks that link microbial ecology with urban planning and management.

